# SEEKING CANDIDATE MOLECULES AS PROGNOSTIC HEALING MARKERS IN CHRONIC VENOUS ULCERS

**DOI:** 10.1101/2020.02.18.954594

**Authors:** Nayara Rodrigues Vieira Cavassan, Noemia Aparecida Partelli Mariani, Caio Cavassan Camargo, Ivan Rodrigo Wolf, Benedito Barraviera, Rui Seabra Ferreira, Guilherme Targino Valente, Erick José Ramos Silva, Hélio Amante Miot, Luciana Patrícia Fernandes Abbade, Lucilene Delazari dos Santos

## Abstract

Seeking and identifying biomarker molecules in inflammatory exudate of chronic venous ulcers (CVUs) can aid health professionals in the healing prognosis. The therapeutic failure or cure is related to the quantitative expression of determinate proteins. This work aimed to identify the proteins expressed in inflammatory exudates from CVUs and correlate them with reduction or increase in the wound size. For 90 days, 28 participants that received standard treatment for 37 CVUs were monitored. The inflammatory exudates were collected before treatment initiation (T=0) and analyzed via the Label-free Shotgun. After 90 days the wound area was reduced in 25 (67.6%) of them. Mass spectrometry analysis of all the inflammatory exudates showed four proteins differentially expressed and related to favorable or unfavorable evolution of the healing process. Complement C3 and ceruloplasmin were identified in all the lesions analyzed and were expressed differentially in lesions that presented diminished area in the studied period. Apoliprotein A1 and neutrophil defensin-1 presented differential expression in ulcers that either did not diminish or augmented their wound area through 90 days. These results suggest that Complement C3, Ceruloplasmin, Apoliprotein A1 and Neutrophil-defensin-1 proteins are potential candidate molecules for prognostic healing markers in chronic venous ulcers.

## INTRODUCTION

Chronic venous ulcers (CVUs) are one of the main public health problems in the Western world [1]. Epidemiological studies performed in the last decade evidenced the frightening perspective of an estimated prevalence between 34.5 and 150.8 million persons affected [2,3-6]. Chronic leg ulcers are lesions that generally last longer than 6 weeks and in some cases more than 20 years, frequently recur and are related to complications from venous insufficiency in the lower limbs [7,8].

The treatment is prolonged, onerous to the health systems, and has important social impact since it contributes to absences from work [9-12]. The patients evolve with restricted mobility, social isolation and a worsening quality of life [13,14]. These aspects affect mental health by increasing sleep disturbances [15], evolving with anxiety and subsequent depression [16]. Given that some cases heal and others do not, science has been searching for years for possible prognostic mechanisms of healing.

In this context, the investigative approach of proteomic analysis of human bodily fluids has been viewed as powerful analytic tool for discovering molecular markers that can aid in the diagnosis and prognosis in diverse pathological conditions [17,18]. Thus, inflammatory exudate from CVUs constitutes a rich biological sample that is easy to collect and, in most cases, neglected by health professionals. This is a complex mixture of proteins, present in the lesion, containing growth factors, matrix proteins, proteinases and cytokines. The variations of these components in the exudate can be utilized as a tool to elucidate the events that modulate the chronicity of these wounds [17,19-21].

Identification of potential biomarkers present in exudate from CVUs can aid in comprehending the processes in the success or failure of healing and in the development of new therapeutic alternatives [19,22], as well as predicting the lesion chronicity prognosis. The present work aimed to identify potential proteins differentially expressed in inflammatory exudate from CVUs, and correlate them with the evolution of the healing process, in order to differentiate healing and no-healing ulcers.

## MATERIAL AND METHODS

### Study Type

Observational study of analytic cohort type.

### Study Population

Thirty-seven (37) ulcers were included from 28 patients aged more than 18 years, attended at the Chronic Ulcers Outpatient Unit of the Botucatu School of Medicine – Universidade Estadual Paulista (UNESP), Botucatu, São Paulo state, Brazil, presenting one or more active CVUs with diameter from two to 15 cm and with evolution greater than six weeks. Patients with ulcers of another etiology, associated with peripheral arterial disease (confirmed through vascular Doppler ultrasound exam (DV 610B, when the ankle-arm index was less than 0.9) or with necrotic tissue were disregarded.

### Study Design

The patients were included consecutively in the order of their arrival at the attending service, in the study in the period from March of 2014 to March of 2015. All the patients were submitted to a clinical-epidemiological questionnaire containing questions relevant to the research and to their knowledge of and agreement to the use of their biological material for academic research obtained by signing Terms of Free and Informed Consent (TCLE) approved by the Institute’s Research Ethics Council (report nº 501.218/2013). No curative or therapy was standardized for this study, with the treatment being prescribed individually according to the indication of the responsible dermatologist. However, the recommended treatments were the standard ones for CVUs, that is, curatives according to the necessities of the wound and compressive therapy. The participants were monitored for 90 days. Exudates were collected from active ulcers for proteomic analysis (Baseline time = T0) and analysis of the ulcer areas (T=0 and T=90).

### Ulcer Area Analysis

The analyses of areas of each ulcer were performed at the moments T=0 and T=90. The drawing of the contours of each ulcer was accomplished on transparent plastic film utilizing a hydrographic pen, and then transferred to a blank sheet of paper. Near the drawing was determined a scale of two centimeters for creation of a photographic reference. Subsequently, photographs were captured of the drawings and references of all the ulcers, which, after being transferred to a computer could be analyzed individually via the software ImageJ (Version 1.52) [23]. Starting from the known reference, the software was calibrated to estimate the area of each lesion from the resolution (pixels/cm) of each ulcer. The ulcer area at T=0 (A0) was utilized as a reference to determine the diminution or lack thereof of the wound at T=90 (A90) by the equation ΔA=A90-A0. The difference between the initial and final areas of each ulcer was denominated the ulcer reduction velocity.

### Collection and Preparation of Material

The collection of exudates and the analysis of the areas were carried out at the moment of the first attendance (T=0), based on the protocol elaborated by Fernandez and collaborators [19]. First, the ulcer was washed with 0.9% saline solution (m/v) of sodium chloride for cleaning, then subsequently dried with sterile gauze. Each lesion was then wrapped with a transparent polyurethane semi-occlusive dressing (Tegaderm®; 3M Health Care, St. Paul, MN, USA) and the patient was maintained at rest for 30 to 60 minutes to wait for natural exudation. The exudate accumulated between the bedding of the ulcer and the dressing was collected with the aid of a disposable-tip micropipette, and then transferred to Eppendorf^®^ tubes, identified and stored in a box containing ice until reaching the laboratory, where they were centrifuged at 14,000g at 4ºC for 10 minutes for sedimentation of all or any cellular debris. The supernatant was stored at −80°C until use.

### Quantification of proteins

The proteins present in the exudate were quantified in triplicate by the method of Bradford [24], (BioRad®; Protein Assay, cod. 500-0001), employing bovine albumin (BSA) as the standard protein. The mean value of each sample was utilized for the quantification calculation. After this procedure the samples were transferred to LoBind-type Eppendorf^®^ tubes and diluted with 0.9% saline solution (m/v) for standardization of their concentration, establishing the relation 50µg/40µL for each sample.

### Digestion of proteins in solution

The samples were digested in individual solution according to the methodology described by Cavassan and collaborators [25], beginning with the steps of reducing and alkylating the proteins. The enzyme trypsin (Promega) was used at the ratio of 1:50 (enzyme: substrate), solubilized in 50 mM ammonium bicarbonate buffer, pH 7.8. Hydrolysis occurred for 18 hours and was interrupted by the addition of 1% (v / v) formic acid in relation to the sample volume. Samples were desalted using Sep-Pak Vac C18 cartridges (Waters) and then lyophilized in SpeedVac ™ (Thermo Scientific) and refrigerated at 4 ° C until the moment of analysis by mass spectrometry.

### Mass Spectrometry Analyses

The analyses by mass spectrometry were accomplished by the methodology by Cavassan and collaborators, described previously [25]. The samples were solubilized in 60 uL of 0.1 % (v/v) formic acid solution; next, a 15 µL aliquot of the tryptic digestion products from each sample were individually injected into a C18 analytical column, 1.7µm BEH 130 (100 µm x 100 mm) in a system of Reverse Phase Liquid Chromatography (RP-UPLC - NanoAcquity UPLC, Waters - Milford, USA) coupled to a Q-Tof PREMIER mass spectrometer (MicroMass/Waters-Milford, USA) for triplicate analyses. The linear gradient used was 2 to 90% (v/v) acetonitrile in 0.1% (v / v) formic acid over 60 minutes at the flow rate of 600 nL/min. The instrument was operated in positive ionization mode and continuous data acquisition was obtained in the molecular mass range from 100 to 2,000 m/z.

### Data Analysis

From the data obtained by mass spectrometry (LC MS-MS), the proteins were identified through the bioinformatics tool Mascot Distiller v.2.3.2.0 (Matrix Science, Boston - USA), utilizing public databases (NCBI, *Homo sapiens* taxonomy, 33,695,097 sequences, available at http://www.ncbi.nlm.nih.gov/protein/?term=homo%20sapiens). Trypsin was employed as proteolytic enzyme, carbamidomethylation as fixed modification (monoisotopic mass of 57.0215Da), oxidation da methionine oxidation as variable modification (monoisotopic mass of 15.9949) and tolerance error of 0.1 Da for the data from MS and MS/MS.

It should be emphasized that the MS/MS data were considered valid according to the statistical algorithm of the MASCOT tool, with identification values *Mascot Scores* above 42 and at least one of the peptide sequences identified with Ion Score values greater than 30. The counting of spectra count for all proteins identified was carried out by the tool Scaffold Q +, and the False Discovery Rate (FDR) was 1% for proteins and 0.1% for peptides, with 95% reliability. Label-free proteins were quantified with the requirement that at least two peptides be in common in the samples.

### SDS-PAGE and Western blotting

The experiments were performed according to the methodology described by Silva and collaborators [26]. Proteins (50µg) were separated by molecular weight on 4-12% (m/v) NuPAGE Bis-Tris gels, at 200 V for 25 min. Proteins were then transferred to polyvinylidene fluoride (PVDF) membranes at 15 V per 1 h. Transfer was verified by Amido Black staining. Membranes were incubated in blocking solution containing 5% milk (m/v) in TBS-T buffer (100 mM Tris-HCl pH 8.0, 150 mM NaCl and 0.05% Tween 20, v/v) for 1.5 h, then incubated with primary antibodies against ceruloplasmin (Abcam, cat. #Ab110449; 0.3 µg/ml), neutrophil-defensin-1 (Abcam, cat. #Ab122884; 2.0 µg/ml) or anti-GADPH (Abcam, cat. #Ab9485; 2.0 µg/ml), used as loading control. After 3x 5-min washed in TBS-T buffer, membranes were incubated with appropriated secondary antibodies conjugated to horseradish peroxidase (Jackson ImmunoResearch, cat. #711-035-152 and #805-035-180; 0.01 µg/ml). Protein bands were detected using SuperSig-nal™ West Femto Maximum Sensitivity Substrate (Thermo Fisher, cat. #34095). Negative control experiments were performed in the absence of primary antibody. Densitometry was performed on the resulting images via the software ImageJ (Version 1.52).

### Interaction network and enrichment analysis

The network of complete interactions of BioGRID databases [27] was obtained through interface of the program Cytoscape [28] limiting the interactions to those present in the specie *Homo sapiens*. In this network, the nodes represent proteins and the edges their physical interactions. To determine the possible biological role of the proteins identified, the selected proteins and their nearest neighbors were extracted from the network and aggregated into a sub-network (subsequently referred to as “subnet”) through the software Igraph [29], implemented in the R statistical analysis [30]. All the nodes of the subnet were submitted to the g:Profiler server [31] to obtain enrichments of the ontology terms referent to biological processes (BP) [32] and the KEGG database pathways [33].

### Statistical Analysis

The spectral counts of each sample were represented from the weighted mean of the triplicates: (3*median+minimum+maximum)/5, with subsequent exclusion of their respective clusters. Proteins identified in less than 30% of the cases were excluded, while for the remainder, the quantifications that resulted in a zero value underwent multiple imputations [34]. Outlier values (>1.5x the interquartile deviation added to the third quartile or subtracted from the first quartile) were win-sorized [35]. Normality of the samples was estimated by the test of Shapiro-Wilk [36].

The independent variables were represented by the identified proteins and the dependent variable by the area reduction of ulcers at T90 (ΔA=A90-A0). The proteins related to the area reduction were identified by the regression technique of Projection of Latent Structures (PLS), with scaled variables. The regression coefficients of the scaled data were compared to the indicator Variables Important in Projection (VIP) by the Volcano plot diagram. Proteins were selected that presented VIP>1 and regression coefficient >0.1 or <-0.1. The data were analyzed in the software packages JMP 10 and SPSS 22. Those proteins that were present in at least 90% of the ulcers studied were considered prognostic molecular marker candidates.

## RESULTS

The present study included 37 CVUs from 28 patients. After calculating the difference between the initial and final areas of the wounds, it could be observed that 25 (67.6%) lesions presented a reduced area at T=90, with a mean cicatrization of 1.68 (±11.76) cm^2^ (Table 1).

**Table 1:**
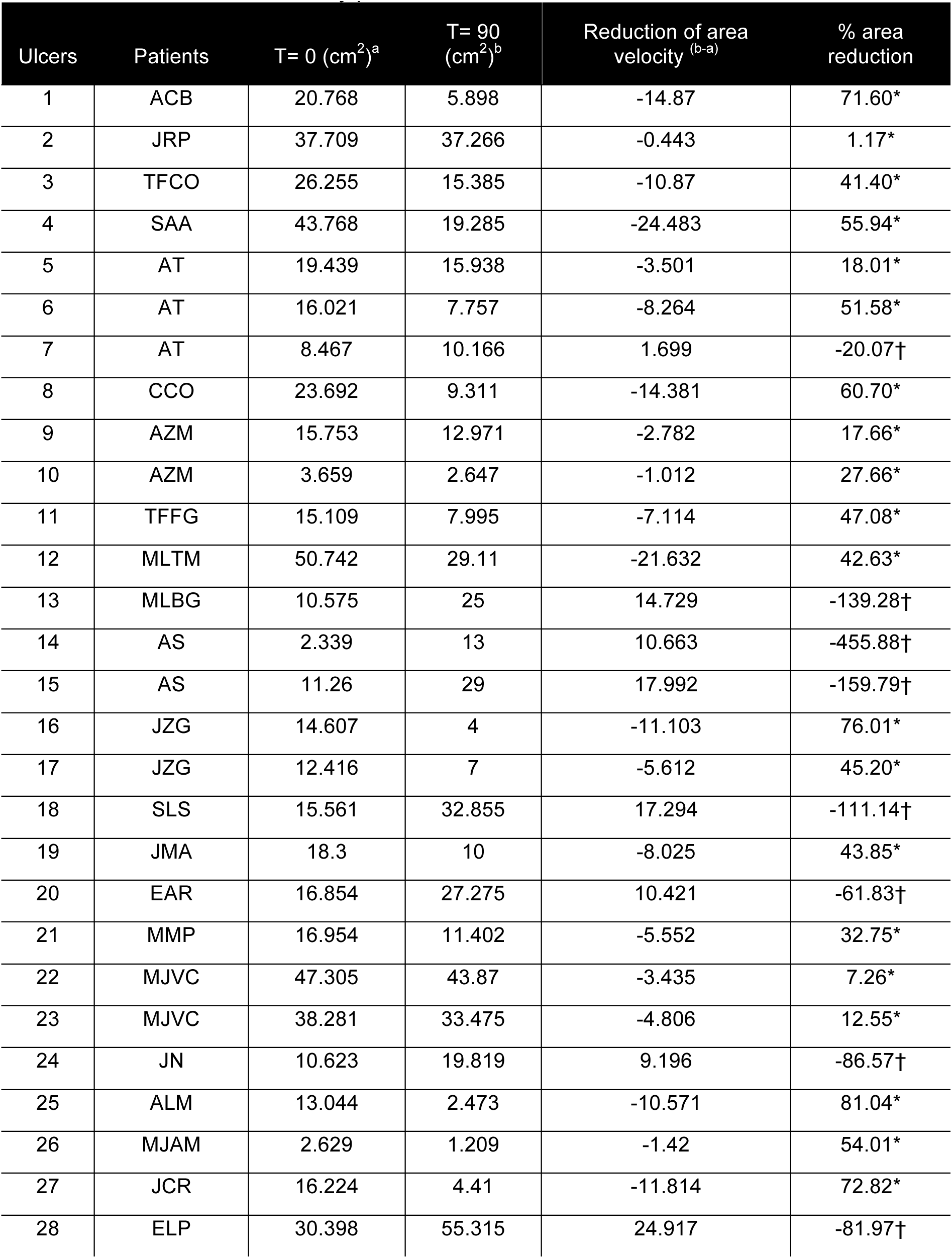
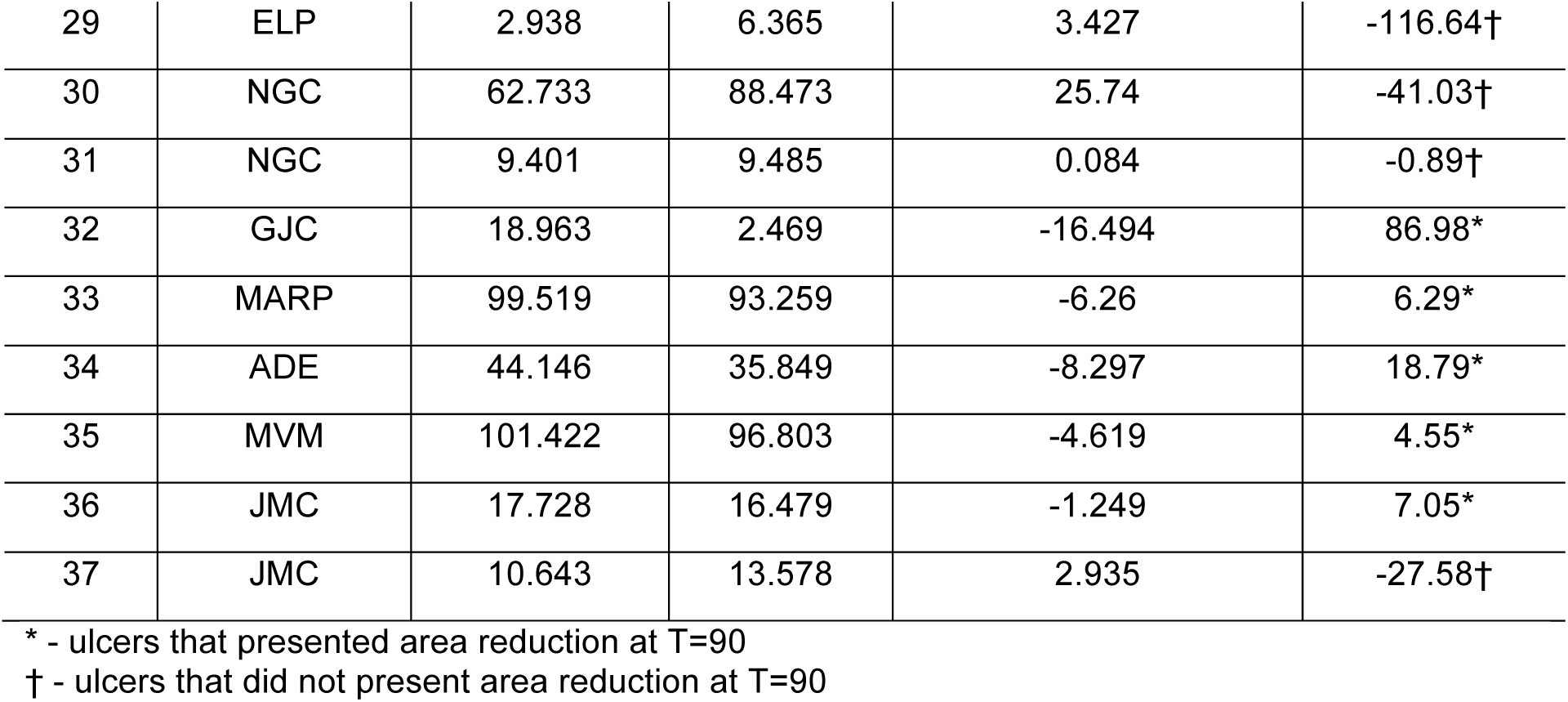
Area of 37 chronic venous ulcers in patients attended at the Outpatient Unit for Venous Ulcers at the UNESP Botucatu Medical School, Botucatu, SP, Brazil, in 2015, which were observed in a 90-day period.

Our recent study by means of *label-free shotgun* mass spectrometry analysis of inflammatory exudates originating from 37 CVUs at T=0, evidenced 76 proteins, which were classified according to their primary role in the healing process [25]. From these data, the correlation of PLS coefficients from 37 scaled samples and from the VIP values indicated four proteins related to cicatrization of CVUs (Figure 1). Multivariate analysis demonstrated that the proteins P4, P8, P17 and P24, respectively identified as Complement C3, Apolipoprotein AI, Ceruloplasmin and Neutrophil-defensin-1, were expressed differently from the others found in the CVU exudates (Table 2) in the presence of ulcers that heal versus those that do not heal.

**Table 2:** Differentially expressed proteins identified in inflammatory exudate das 37 chronic venous ulcers treated in Chronic Ulcers Outpatient from Botucatu Medical School, Sao Paulo – Brazil, in 2015.

| Protein | Name | Acess code | Function | Expression<br>* |
| --- | --- | --- | --- | --- |
| P4 | Complement C3 | P01024.2 | Immunomodulation | H |
| P8 | Apolipoprotein A-I | P02647.1 | Transporter protein | NH |
| P17 | Ceruloplasmin | P00450.1 | Transporter protein | H |
| P24 | Neutrophil-defensin-1 | P59665.1 | Antimicrobial activity | NH |

\* H: differential expression in CVUs that had area reduction; NH: differential expression in CVUs that did not present reduction.

**Figure 1:**
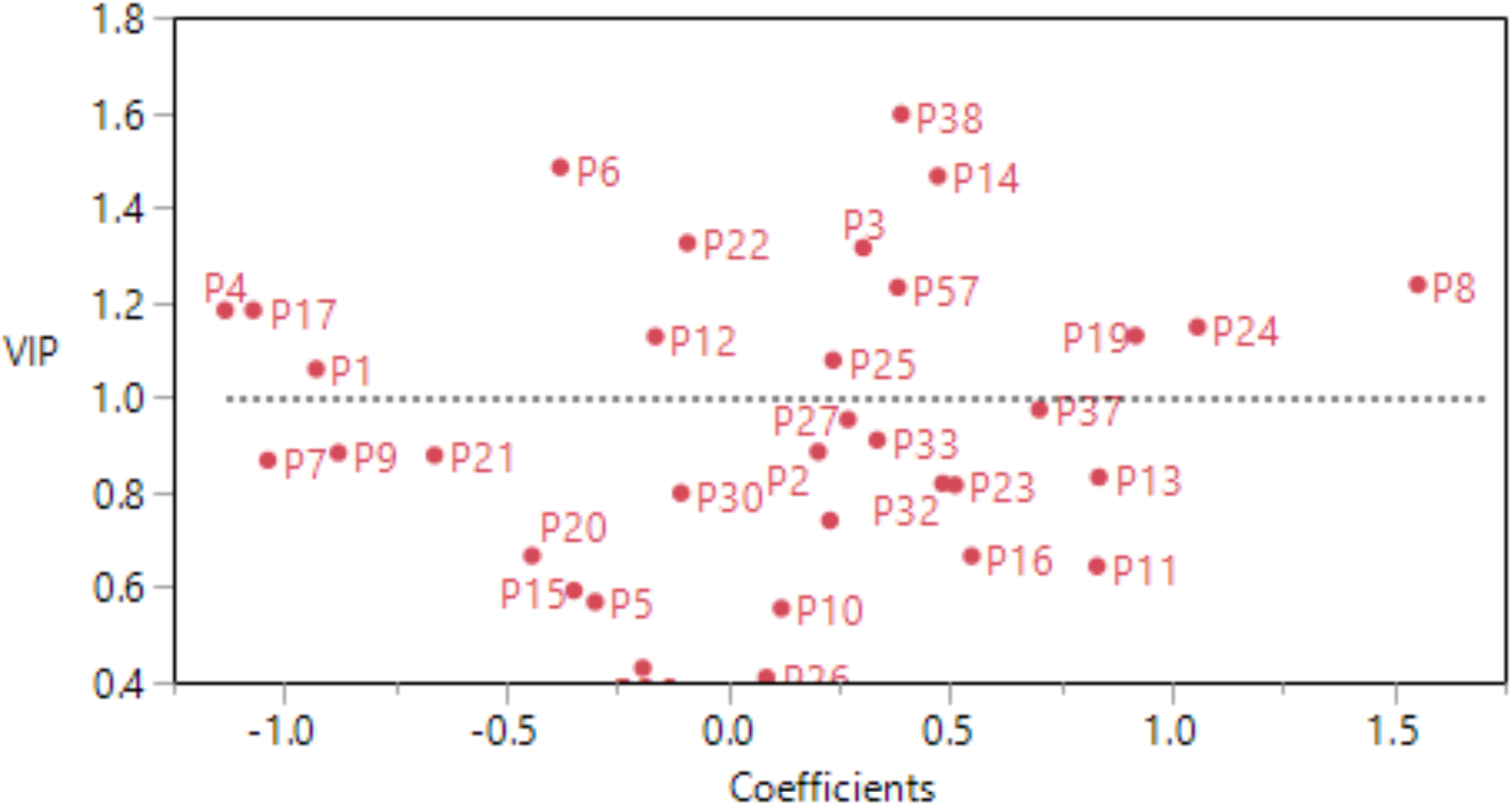
Diagram (Volcano plot) of proteins with differential expression identified among CVUs obtained by multivariate statistical analysis. Complement C3 (P4) and ceruloplasmina (P17) presented differential expression in CVUs that showed reduced area (CoefT90 negative) and ApoAI (P8) and HP1 (P24) in CVUs that did not present this outcome (CoefT90 positive).

Despite the lack of consensus as to the minimum prevalence values, we believe that these molecules should be present in at least 90% of the cases studied, thus reflecting the population as a whole and presenting a highly predictive value as to the question analyzed. The proteins Complement C3 and Ceruloplasmin were identified in all the lesions analyzed and were expressed differentially in lesions that presented diminution from their initial area to the end of the study. The proteins Apoliprotein A1 and Neutrophil-defensin-1 presented differential expression in ulcers that did not heal or increased their initial area across the 90 days. The identification data of these proteins are displayed in Table 2.

In contrast to the evidence indicated by mass spectrometry analyses, western blotting analysis corroborated an increase in the expression of Ceruloplasmin and a decrease in the expression of Neutrofin-defensin-1 in exudates from healing ulcers in comparison to non-healing ulcers. Western blotting experiments confirmed the presence of Ceruloplasmin as a band with apparent molecular mass of 144 kDa in exudates from both healing and non-healing ulcers (Figure 2A). Likewise, the expression of Neutrofin-defensin-1 was also detected in all exudates, but as a double-band with apparent molecular masses of 7 and 5 kDa, which are likely to represent its full-length and secreted isoforms (Figure 2A). No bands were detected in experiments performed in the absence of primary antibodies (data not shown).

**Figure 2.**
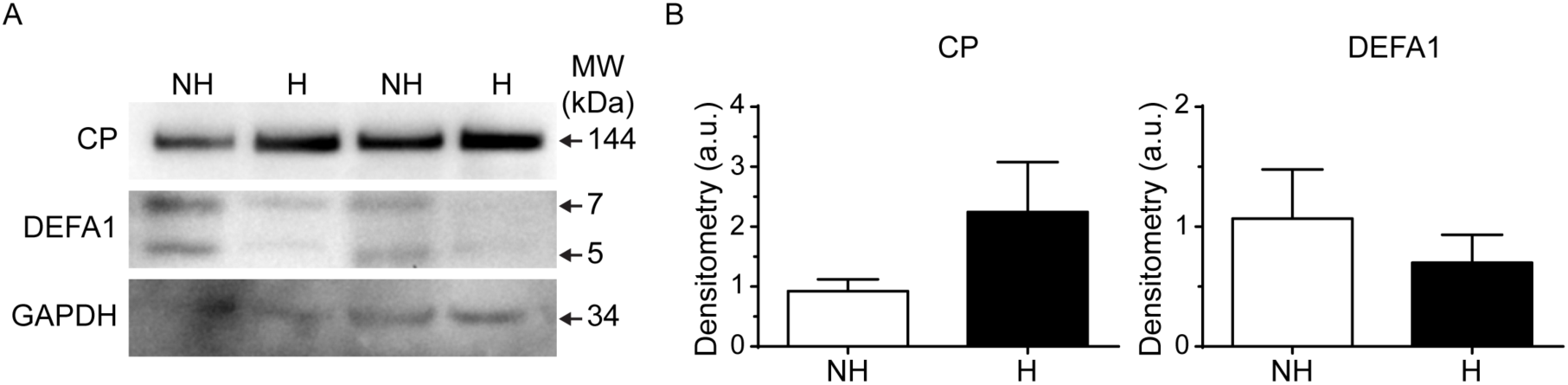
Expression of ceruloplasmin (CP) and α-defensin-1 (DEFA1) in exudates from non-healing ulcers and healing ulcers. **(2A)** Representative CP and DEFA1 detection by Western blot using exudates from non-healing (NH) ulcers and healing (H) ulcers as indicated. Arrows indicate apparent molecular masses (kDa) for CP (144 Da), DEFA1 (7 and 5 Da), and GAPDH (34 Da), used as internal control. **(2B)** Densitometric analysis of the Western blot results. Each sample was normalized to its respective internal control and expressed as levels relative to HH samples. Densitometric analysis for DEFA1 was performed using the lower band. Results are expressed as mean ± SEM of experiments performed with samples from 3-4 exudates/group.

The abundance of the bands corresponding to Ceruloplasmin and Neutrofin-defensin-1 were differentially affected when exudates from healing ulcers and non-healing ulcers were compared by densitometry analysis. We observed a ~2.2-fold increase and a ~0.4-fold decrease in the abundance of Ceruloplasmin and Neutro-fin-defensin-1 in exudates from healing ulcers in comparison to non-healing ulcers, respectively (Figure 2B).

The network of protein-protein interactions of proteins differentially expressed in inflammatory exudate from venous ulcers was proposed in order to elucidate the correlation existent between these candidates for prognostic markers (Figure 3) in reference to cicatrization and initial area across the 90-day evaluation period. The enrichment of ontological terms revealed that among the pathways identified are “*Staphylococcus aureus* infection”, “Complement and coagulation cascades”, “Cholesterol metabolism”, “Pertussis and Fat digestion and absorption”, and among the GOs terms are “high-density lipoprotein particle remodeling”, “response to stress”, “plasma lipoprotein particle organization”, “plasma lipoprotein particle remodeling and protein-lipid complex remodeling” (Supplementary Material S1).

**Figure 3:**
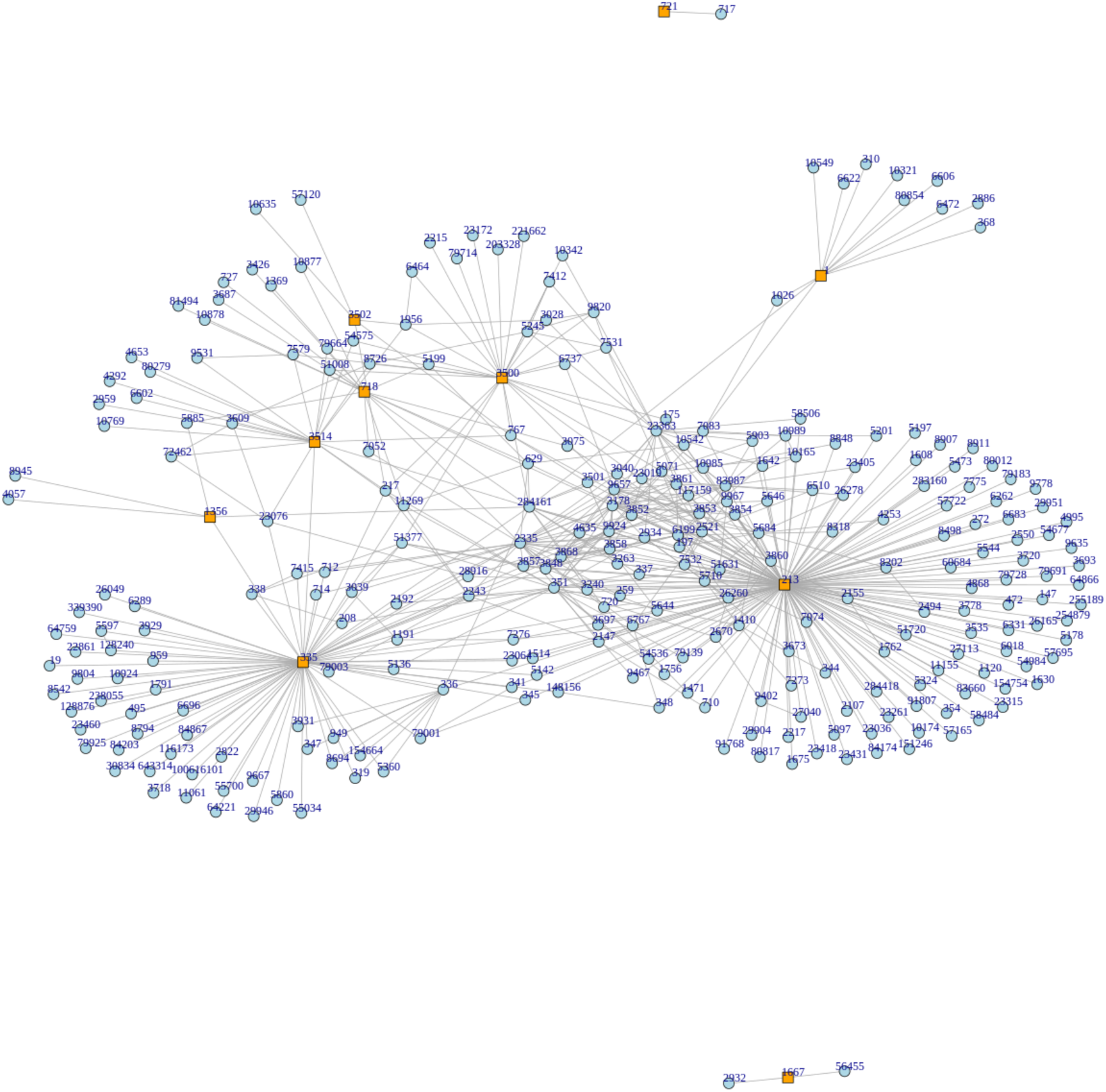
Network of interactions among differentially expressed proteins in inflammatory exudate from chronic venous ulcers that had areas reduced and those not reduced in a period of 90 days. Each vertex/node represents a protein while each edge represents an interaction between the vertices. The nodes 718 and 335 represent the proteins Complement C3 (P4) and Apoliprotein A1 (P8).

## DISCUSSION

Chronic wounds have an elevated financial impact on the health systems of the Western world, especially in relation to the quality of life and absences from work among the affected patients [1]. Although CVUs represent a significant clinical challenge, Australia and other developed nations have prioritized investments in research and development rather than increasing the number of hospital beds for the treatment of these patients [37].

Proteomic analysis of leg ulcer exudate has been indicated as one of the main approaches for investigating molecules with differential expression, suggesting them as potential biomarkers for the healing of these lesions [1,38,39]. In 2017, Broszczak and collaborators [1] indicated that spatial-temporal analyses of wounds, together with the integration of multiple Omic datasets, could provide information on the main candidate molecules that lead to wound chronicity.

Our group has achieved the pioneering first step towards identifying seventy-six proteins from inflammatory exudates from CVU patients, classifying them according to their primary role in the cicatrization process. Thus, we can correlate clinical and epidemiological data with the expression of proteins [25]. From these observations, the present study permitted us to indicate four proteins differentially expressed and present in CVU inflammatory exudates as potential candidates for healing prognosis, namely: Complement C3, Apolipoprotein AI, Ceruloplasmin and Neutrophil-defensin-1.

It is known that the protein Complement C3 is one of the molecules that compose the complement system, being the main modulator of inflammatory response when tissue damage occurs. It participates in the activation of the innate immune system and is responsible for acting in the process of recognizing and clearing pathogens [40-43]. The activation of the complement system occurs by hydrolysis of the protein Complement C3, releasing the fragments C3a and C3b. In turn, these fragments are deposited on the membranes of damaged cells, triggering a cascade of events such as the opsonization and attack complex formation, responsible for inducing inflammation and, subsequently, the elimination of injured cells. Cicatrization dysfunctions may occur when there is a loss of complement system tolerance to C3 fragments, thus inducing autoimmune mechanisms, chronicity and tissue damage [40,41]. The immune response that takes place in the CVU microenvironment induces a series of enzymatic reactions, which can culminate in a normal inflammatory response or in tissue inflammation that is unsatisfactory and uncontrolled [44].

A favorable inflammatory response results in tissue progression to the phases of granulation and remodeling of the wounds. However, some studies suggest that there is an increase of proteolytic activity in the microenvironment of CVUs generating the degradation of complement system compounds and impeding lesion healing by stagnation in the inflammatory phase [45-47].

Furthermore, CVUs are susceptible to the action of a wide variety of bacterial agents since they disrupt the first line of defense of the innate immune system by breaking the physical barrier of the skin [48-51]. The presence of microorganisms activates the complement system through the interaction of alternative pathways and the protein lectin. However, the identification and opsonization of infectious agents are dependent on the binding of the protein lectin with mannose (MBL) [52-54]. Some studies suggest that a deficiency in serum MBL levels in patients with CVUs culminates in the incapacity of complement system activation, facilitating the installation of infections [55,56]. Thus, the differential expression of protein Complement C3 in the lesion microenvironment is the result of the activation of the defense system against infectious agents, and also suggests the installation of the inflammatory process in the region. The overexpression of this protein may be related to the speed and efficacy in the formation of the antigen attack complex, regarding the equilibrium of serum MBL levels, with consequent diminution of the local bacterial load, tissue restoration and favorable progression of the healing process.

Yet the protein Ceruloplasmin is the main carrier protein of plasmatic copper. It binds to immune system cells, presents enzymatic and anti-inflammatory activity, and has its expression augmented in response to hypoxia, triggering an increase of thrombin and pro-inflammatory cytokines in the blood [57-62]. Its over-expression in the lesion microenvironment may facilitate healing by its ability to bind with the protein lactoferrin, favoring Fe3 + sequestration from inflammatory fluid, essential for bacterial growth. Thus, the decrease or absence of iron ions in the wound protects the wound against the action of pathogenic bacteria, besides inhibiting the production of the enzyme myeloperoxidase with consequent reduction of oxidative stress [63-65].

Kostevich and collaborators [66] suggest that the elevated proteolytic and oxidative action during an inflammatory condition may result in structural transformations of Ceruloplasmin, augmenting its interaction with molecules, including macrophage inhibition factor (MIF), favoring pro-inflammatory activity in the tissues. The same was observed by Laura Anca and collaborators [67], who described a positive association between oxidative stress, insulin resistance and inflammation, and elevated levels of ceruloplasmin in obese children. Although Ceruloplasmin appears to perform both pro-and anti-inflammatory roles, its function in inflammatory conditions still requires new investigations [68]. Thus the presence of proteolytic and oxidative activity at the lesion site appears to be associated with the structural integrity of the protein Ceruloplasmin, denoting the beneficial effect of differential expression of this molecule in the healing of CVUs.

It is know that leg ulcers are in many cases associated with the presence of bacteria and fungi. These microorganisms can affect cicatrization by producing toxins and biofilms, creating a physiochemical barrier against the immune system of the host [69,70]. In this context, some proteins found in the current study appear to perform antimicrobial activity against a variety of microorganisms. The protein *Neutrophil defensin 1* is a molecule from the family of defensins stored in granules of neutrophils, and performs the function of increasing the permeability of bacterial cells. Nevertheless, these antimicrobial factors are not specific to the pathogens. If a continuous dysregulated stimulus of neutrophils occurs at the wound site, tissue growth factors are degraded and consequently, wound regenerated is impaired [71].

Lundqvist and collaborators [72] corroborate our findings, indicating that differentiated expression of Neutrophil defensin-1 in exudate and in tissue of CVUs occurs in response to infectious agents. This protein can exert pro-inflammatory and cytotoxic effects, thus contributing to elevated inflammation – a characteristic of CVUs that do not heal. Furthermore, Cardot-Martin and collaborators [73] demonstrated that the defensins produced by neutrophils neutralize bacterial cytotoxins produced by *Staphylococcus aureus*, which is very common in CVUs.

Yet Apoliproteins of the types AI and AII (APOAI and APOAII) form the compound HDL (*High density lipoprotein*), upon binding to lipids. It is believed that when free of binding with HDL, they combine with free particles of VLDL (*Very Low Density Lipoprotein*) compounds, which are considered “bad cholesterols”, thus completing their reverse transport and performing a protective function against atherosclerosis [74]. Li and collaborators [75] reported an increase of atherosclerosis cases in an aging population, suggesting that the elevation of atherosclerosis in the elderly may be related to diminished levels of APOs in carrying plasmatic VLDL compounds, thereby inducing greater VLDL deposition in the walls of veins and arteries [76].

Known for neutralizing products released by gram-positive (lipotechoic acid) and gram-negative (lipopolysaccharide) bacterial cells, HDL participates in neutralization of inflammatory processes mediated by macrophages [77]. Thus, the presence of APOAI in CVUs might indicate an attempt by the organism to contain high levels of colonization and infection, highly present in this type of lesion, and diminish their potential to heal. Low serum APOA levels were described as indicative of a poor prognosis in sepsis, for forming few HDL complexes [77].

## CONCLUSION

Given our results and the comprehension of the biological and molecular roles, it is suggested that the proteins Complement C3, Ceruloplasmin, Apoliprotein A1 and Neutrophil-defensin-1 present in the inflammatory exudates of CVUs are potential candidates for prognostic markers of cicatrization. Multi-centric clinical trials are necessary to validate these findings. If they are confirmed, it would be desirable to develop rapid diagnostic/prognostic kits to assist the healthcare professional in the primary care of these patients. This would not only optimize financial resources and permit a better treatment choice, but also alert the patients as to the possible evolution of their disease.

## Availability of data and material

Spectral data from LC-MS/MS have been uploaded to the Atlas Peptides Repository from Institute for System Biology (http://www.peptideatlas.org/) with the dataset identifier PASS01190.

## ACKNOWLEDGEMENTS

Special thanks are extended to the Center for the Study of Venoms and Venomous Animals (CEVAP) of São Paulo State University (UNESP), the Experimental Re-search Unit (Unidade de Pesquisa Experimental - UNIPEx) and the Chronic Ulcers Outpatient Unit of the Dermatology Service from of the Botucatu Medical School (Faculdade de Medicina de Botucatu, FMB) for making available all the resources and equipment that contributed to this work. We acknowledge the Mass Spectrom-etry Laboratory at the Brazilian Biosciences National Laboratory, CNPEM, Campi-nas, SP, Brazil, for its support with the mass spectrometric analysis.

